# Dissociation of dorso-lateral and dorso-medial prefrontal cortex contributions to familiarity and recollective processes in primates

**DOI:** 10.1101/2020.01.16.909812

**Authors:** Zhemeng Wu, Martina Kavanova, Lydia Hickman, Fiona Lin, Erica Boschin, Juan M. Galeazzi, Lennart Verhagen, Mark J. Buckley

## Abstract

According to dual-process theories, recognition memory draws upon both familiarity and recollection. It remains unclear how primate prefrontal cortex (PFC) contributes to familiarity and recollection processes but frequency-specific neuronal activities are considered to play a key role. Here, non-human primate (NHP) electrophysiological local field potential (LFP) recordings first showed that a specific subregion of macaque PFC (i.e., dorsolateral PFC, dlPFC) was implicated in task performance at a specific frequency (i.e., increased beta power in the 10-15 Hz range observed in correct versus error trials) in a specific phase of a recognition memory task (i.e., during sample presentation). Then, to assess generalization to humans and causality we targeted left human dlPFC (BA 9/46) as well as left dorsomedial prefrontal cortex (BA 8/9) for comparison, and also vertex as a control, with transcranial magnetic stimulation at a frequency in the middle of the low-beta range observed in NHP (i.e. 12.5 Hz) and compared that to non-frequency-specific stimulation, and also to a no-stimulation control, during occasional sample presentations within a similar task. Hence we investigated hypotheses about the causal importance for human memory of a location-specific, frequency-specific, and task-epoch-specific intervention derived directly from the NHP electrophysiological observations. Using a dual-process signal detection (DPSD) model based on analysing receiver operating characteristics (ROC) curves, we showed beta-frequency TMS caused decreased recollection when targeted to human dlPFC, but enhanced familiarity when targeted to dorsomedial prefrontal cortex. Non-frequency-specific patterns of stimulation to all sites, and beta-frequency stimulation to vertex, were all without behavioural effect. This study provides causal evidence that PFC-mediated contributions to object recognition memory are modulated by beta-frequency activity; more broadly it provides translational evidence bridging NHPs and humans by emphasizing functional roles of beta-frequency activity in homologous brain regions in recognition memory.

**Highlights:**

- low beta power in NHP dlPFC during stimulus encoding was related to behaviour
- human rTMS study used parameters derived from NHP observations to test causality
- low beta rTMS to human dlPFC, but not dmPFC, impairs recollection
- low beta rTMS to human dmPFC, but not dlPFC, enhances familiarity
- provides cross-species validation of prefrontal beta power to primate recognition

## 1. Introduction

There is also a long-standing literature on the necessity of PFC for supporting object recognition memory based on lesion studies in macaques (Bachevalier and Mishkin, 1986; Kowalska et al., 1991; Levy and Goldman-Rakic, 1999; Meunier et al., 1997; Mishkin and Manning, 1978; Parker, 1998; Petrides, 2000, 1995). Early neuronal recording studies in macaque prefrontal cortex (PFC) suggested sustained neuronal spiking might encode stimuli across delays in object recognition memory tasks (Fuster and Alexander, 1971; Rao et al., 1997; Wilson et al., 1993). More recently other mnemonic mechanisms for this kind of working memory task in PFC have since been proposed (Lundqvist et al., 2016; Stokes, 2015). Consideration as to the role of neuronal synchrony (e.g. within and between-area oscillatory coherence), including at different frequencies, and their importance in helping mediate cognition including learning and memory has also become a key area of systems neuroscience research (Benn et al., 2016; Buzsáki and Schomburg, 2015; Fell and Axmacher, 2011; Fries, 2015; Gregoriou et al., 2009; Herweg et al., 2016; Köster et al., 2014; Pipa, 2009; Womelsdorf et al., 2007).

In this context the local field potential (LFP) electrophysiological signal is thought to reflect a summation of local transmembrane currents (Buzsáki et al., 2012) such that LFP amplitude may reflect local synchrony within an area. One influential theory is that neuronal synchrony also facilitates inter-area neuronal communication; for instance, given rhythmic inhibition within local networks, two neuronal ensembles may have a greater influence on each other when their temporal interaction windows open at the same times (i.e. rhythmic synchronization *within* areas also synchronized *between* areas) (Womelsdorf et al., 2007). Relatively few studies to-date have analysed LFP activity in non-human primate (NHP) PFC during working memory tasks. One recent electrophysiological LFP recording study in NHPs indicated discrete and dynamic modulation of beta (20-35 Hz) and gamma oscillations (45-100 Hz) in lateral PFC during encoding period of working memory, which suggests beta oscillations in PFC reflect a default network state while are interrupted by gamma oscillations when encoding or decoding the stimuli (Lundqvist et al., 2016). Kornblith et al. (2016) have found that beta synchrony (16-30 Hz) in lateral PFC increases (although beta power decreases) during multiple stimulus presentation while decreases (although beta power increases) during memory delay; moreover, both gamma synchrony and power (50-100 Hz) in lateral PFC increase during multiple stimulus presentation. The results of this study indicate that roles of low-frequency oscillations in top-down processing and higher-frequency oscillations in bottom-up processing (Kornblith et al., 2016). Another electrophysiological study shows that small groups of neurons in macaques probably comprising a cortical column participate in high-gamma oscillations around 80 Hz and their activity carries partial information about the memorized stimulus, while larger networks in and most likely beyond PFC appear to be coordinated by coherent low-gamma- and beta-oscillations (14-45 Hz) which are correlated with performance but not stimulus content (Pipa, 2009). Although studies have analysed LFP activity in macaque PFC during recognition memory they facilitate comparison between PFC regions and even cortical layers (Bastos et al., 2018). Moreover, in humans, a significant body of research maintains that human recognition memory draws upon two processes: familiarity and recollection (Wixted, 2007; Yonelinas, 2002, 2001, 1994; Yonelinas et al., 2002, 1998; Yonelinas and Parks, 2007). One influential neuropsychology review concluded that PFC patients were numerically worse than controls at recognition but deficits in recall were more profound (Tulving et al., 1995). However, there is no consensus as to whether different sub-areas of PFC contribute differently to familiarity and recollection. Using paradigms that aim to dissociate recollection from familiarity (e.g. receiving operating characteristic (ROC) curve plotting across confidence levels, or dissociations of subjective reports of ‘remembering’ versus ‘knowing’), human neuropsychological investigations present conflicting results. In some, lateral PFC lesions cause deficits in familiarity rather than recollection (Aly et al., 2011; Duarte et al., 2005; MacPherson et al., 2008), whereas others (Wheeler and Stuss, 2003) report that dorsolateral PFC (dlPFC) lesions don’t impair either, in contrast to frontopolar lesions which only impaired recollection. Evidence from human neuroimaging studies also provides mixed evidence as to the relative contributions of PFC sub-regions to familiarity/recollection (R. N.A. Henson et al., 1999; Richard N.A. Henson et al., 1999; Horner et al., 2015; Johnson et al., 2013; Kafkas and Montaldi, 2012; Skinner and Fernandes, 2007; Solstad et al., 2006). A recent review of previous neuroimaging studies involving human PFC similarly reports a mixture of involvement in recollection/familiarity in different parts of PFC (Scalici et al., 2017), which suggests a complicated characterization of PFC in association with these component memory processes supporting recognition memory. In a combined fMRI-EEG study in humans, an alpha-theta synchronization (4-13 Hz) during recollection is associated with increased functional connectivity between the PFC and hippocampus, indicating low-frequency oscillations support the PFC-hippocampal network dynamics during recollection process (Herweg et al., 2016). Another human electroencephalography study shows that a coupling between prefrontal theta and parietal gamma oscillations occur during successful memory retrieval, which suggests an essential role of an interaction between low- and high-frequency oscillations in memory retrieval (Köster et al., 2014).

The homologies between NHP PFC and human PFC, as evidenced by long-standing comparative cytoarchitectural analyses (e.g. Petrides and Pandya, 2007) and from more recent comparative functional connectivity profiles (Neubert et al., 2015), help confirm NHPs as a good animal model to investigate the neural activities and synchronies underlying recognition memory although the debate over whether animals can ‘recollect’ per se is still ongoing. Nonetheless, the neural signals (e.g. finer grained information on LFPs) available in NHPs that are recorded direct from electrodes within specific brain regions may provide important information about the neural mechanisms that may give rise to recollection and familiarity in humans.

In this study we aimed to investigate the causal role of subregions of human PFC in recollection and familiarity using non-invasive transcranial magnetic stimulation (TMS). TMS can be used to intervene in on-going neural processing by administering either single pulse or several repetitive pulses (rTMS) either before or during task performance to modulate ongoing neural oscillations (Huang et al., 2005; Rossi and Rossini, 2004). It has previously been observed that behavioural performance in humans can be either disrupted or enhanced by adopting TMS in a frequency-specific manner (Albouy et al., 2017; Chanes et al., 2013; Elkin-Frankston et al., 2011; Thut et al., 2011). For example, perceptual discrimination has been enhanced by applying TMS over a frontal eye field (an area of human PFC) at high-beta frequency, while response criterion has been lowered by delivery of TMS over the same region at gamma frequency (Chanes et al., 2013). In order to generate narrower hypotheses about causality as to which specific PFC area might be targeted and at what point in the working memory task and at what frequency we sought insight from a simple preliminary investigation in a single NHP that we previously trained to perform a similar task and from which we recorded LFP activity from a multi-electrode array implanted in dlPFC (additional animals were not available as the NHP data was a pilot/parameter defining stage for a subsequent larger-scale longer-term NHP project still in progress).

In the present NHP observations, we found a strong correspondence between increased beta LFP power from electrodes in dlPFC around the time of sample presentation onset in our object recognition memory paradigm and memory performance; the significant sample phase activity was transient, lasting no more than 200 ms and with a frequency range of 10-15 Hz. It’s relation to performance was determined by confirming that the power was significantly higher in trials than proceeded to be correct trials (in the subsequent choice phase) than in trials that proceeded to be error trials. The desynchronization hypothesis (Simon Hanslmayr et al., 2012; Holmes et al., 2018) suggests that PFC neurons desynchronize to retain stable memories. We observed robust beta power that was greater in correct trials but which didn’t persist throughout the entire sample presentation; therefore it may be hypothesized accordingly that transient robust beta power coupled with subsequent efficient beta power desynchronization supports effective memory encoding. To assess both causality and generalizability across primate species we hypothesized that targeting the homologous human region with beta TMS in the same low beta frequency range observed (i.e. our TMS stimulation frequency was chosen as 12.5 Hz being right in the middle of beta range observed in NHP dlPFC) throughout the entire sample presentation in a similar task may lessen desynchronization and disrupt effective memory encoding for those samples. As a control for this hypothesis we included a randomly timed stimulation condition in other blocks (same number of pulse, same duration, but with no inherent frequency) as this non-beta (random) frequency stimulation was presumed to be less likely to prevent beta desynchronization. As human activation peaks related to familiarity were previously generally found to be located more caudally, corresponding to dorsomedial PFC (dmPFC), in meta-analyses (Scalici et al., 2017), we further hypothesized that targeting dlPFC would primarily affect recollection indices, while targeting dmPFC (our chosen comparison target for this reason) would primarily affect familiarity indices. We included vertex as a control region wherein we hypothesized no effect of TMS by either of the two stimulation protocols.

## 2. Materials and Methods

### 2.1 Experiment 1: NHP electrophysiological study

#### 2.1.1 Subject

Neural data was recorded in one young adult female macaque monkey (*Macaca mulatta, age 8 years, weight 10-13 kg*). All animals in our lab are socially housed (or socially housed for as long as possible if later precluded, for example, by repeated fighting with cage-mates despite multiple regrouping attempts) and all are housed in enriched environments (e.g. swings and ropes and objects, all within large pens with multiple wooden ledges at many levels) with a 12hr light/dark cycle. The NHP always had *ad libitum* water access 7 days/week. Most of its daily food ration of wet mash and fruit and nuts and other treats was delivered in the automated testing/lunch-box at the end of each behavioral session (this provided ‘jack-pot’ motivation for quickly completing successful session performance; supplemented by trial-by-trial rewards for correct choices in the form of drops of smoothie delivered via a sipping tube) and this was supplemented with fruit and foraging mix in the home enclosure. All animal training, array implantation surgery, and experimental procedures were performed in accordance with the guidelines of the UK Animals (Scientific Procedures) Act of 1986, licensed by the UK Home Office, and approved by Oxford’s Committee on Animal Care and Ethical Review.

#### 2.1.2 Surgical procedures

After the basic behavioural and initial task training phase was complete the animal first received a titanium head-post, implanted posteriorly and secured to the cranium with titanium cranial screws through the legs of the post (the legs of the head-post were pre-shaped to fit the precise morphology of the skull in the region according to a 3D printed skull model based on pre-operative structural MRI scans); the reflected skin and galea was sutured and the wound closed around the base of the head-post.

Later, after more behavioural training with head-fixation was complete and task-performance satisfactory, the NHP received surgical implantation of microelectrode arrays (Utah arrays, Blackrock Microsystems). A bone flap was raised over the left anterior prefrontal cortex, the dura mater was cut and reflected, and Utah arrays implanted directly into the cortex; the dura mater was sewn back over the arrays where possible and the bone flap was replaced. Two reference wires connecting to the pedestals were left under the dura away from the site of the array and these and the wire bundle connecting to each electrode in the Utah array exited the cranium through a gap at the edge of the craniotomy and ran from there to the pedestal which was secured to the cranium (located away from the edge of the craniotomy) with titanium cranial screws through its base; to complete the procedure the wound was then closed in layers. A pedestal cap was screwed on to the top of the pedestal to protect the gold contacts (the cap was subsequently removed each day, while the NHP in the recording session, to connect the digital head-stage direct to the pedestal’s connectors at which point recording commenced).

The operations to implant the head-post and later to implant the microelectrode arrays were performed in aseptic conditions with the aid of an operating microscope for intra-cranial surgeries. The monkeys were first sedated with ketamine (5 mg/kg), medetomidine (20mcg/kg) and midazolam (0.1 mg/kg) given i.m., intubated, and then artificially respirated using a mixture of carrier gases (oxygen/medical air) and volatile anaesthetic. Surgical depth of anaesthesia was maintained throughout the surgery with sevoflurane (1.0-2.0% to effect) and injectable adjuncts (fentanyl 5 mcg/kg/hr i.v., dexmedetomidine 0.5 mcg/kg/hr i.v.). On average three doses of steroids (methylprednisolone 20 mg/kg i.v. every 4-6 hrs) and in the case of intra-cranial surgery a bolus of mannitol (1000 mg/kg i.v.) were given on the day of surgery to protect against intraoperative brain edema and postoperative inflammation. Steroids were continued in the postoperative phase (dexamethasone 0.2 mg/kg s.c., daily for 5 days). The animals were given an antibiotic (30 mg/kg of amoxicillin intraoperatively every 2 hours, and 17.5 mg/kg daily postoperatively) for prophylaxis of infection. Additional intraoperative medication included atropine (10 mcg/kg i.v. as required), an H2 receptor antagonist ranitidine (1 mg/kg i.v.), meloxicam (0.2 mg/kg i.v.) and crystalloid fluids (Hartmann’s solution 3-5 ml/kg/hr). Heart rate, oxygen saturation of hemoglobin, mean arterial blood pressure, end-tidal CO2, body temperature, and respiration rate were monitored continuously throughout surgery. Postoperative analgesia was provided via opioids (methadone 0.3 mg/kg i.m. or buprenorphine 10 mcg/kg i.m.) and nonsteroidal anti-inflammatory agents (0.1 mg/kg of meloxicam, p.o./i.m., 10 mg/kg acetaminophen p.o.). A proton pump inhibitor (Omeprazole 0.5 mg/kg) was given daily to protect against gastric ulceration as a side effect of the combination of steroid and nonsteroidal anti-inflammatory treatment.

This study records data from one 32-electrode Utah array inserted into dlPFC (area 9/46, just dorsal to the principal sulcus, see Fig. 1B); the electrodes were arrange on a 6 × 6 grid embedded in silicone; the electrode length was 1.0 mm with interelectrode spacing of 0.4 mm. All the microelectrodes were made of platinum (Pt).

**Figure 1.**
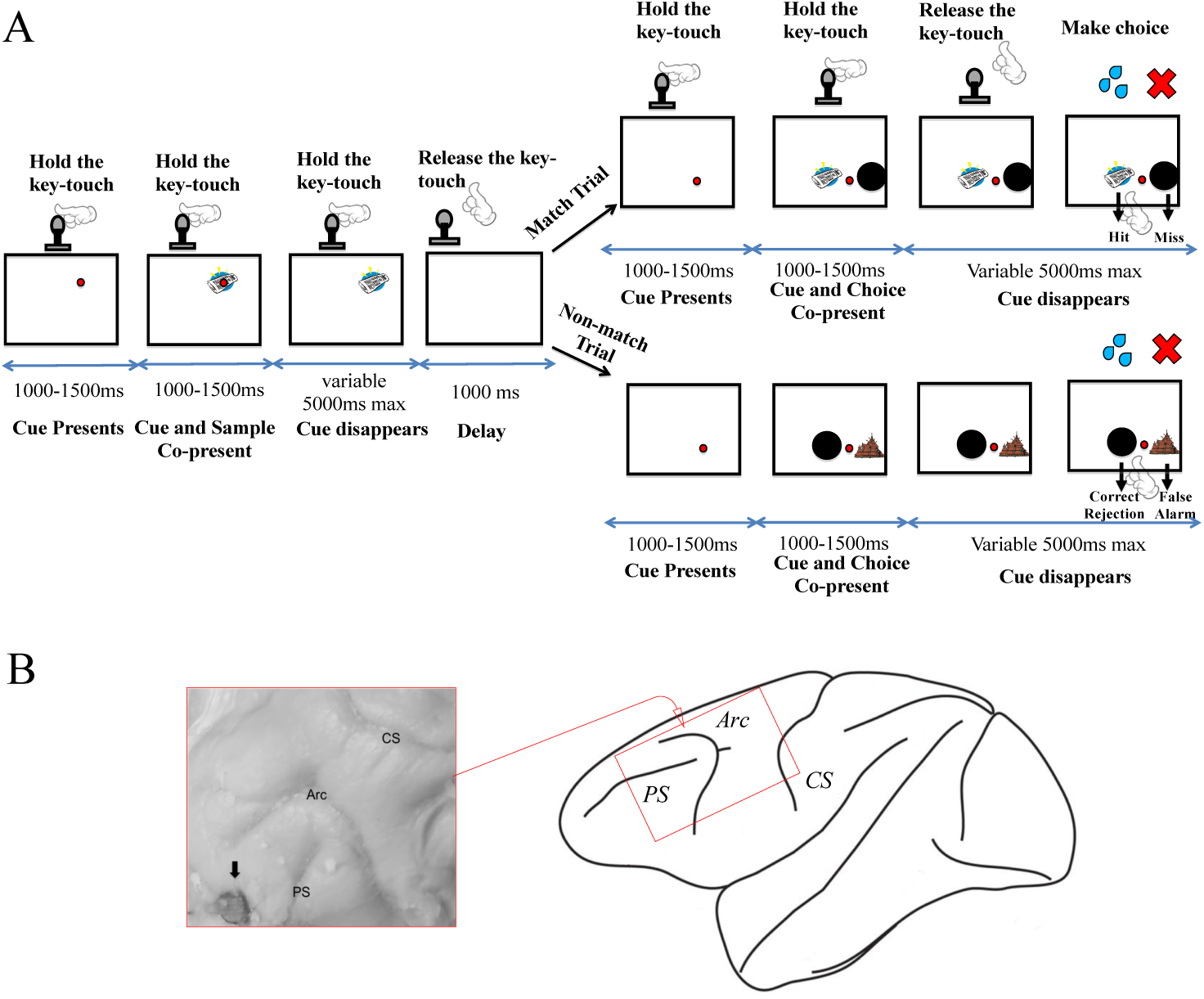
NHP recognition memory task structure and recording sites from dlPFC array in in NHP. Panel A depicts one trial in the electrophysiological study. The NHP initiated the task by holding the key-touch device once the red dot cue appeared through a variable time of 1000-1500 ms, after which a sample object image was shown on the screen behind the red dot cue (encoding phase). The NHP was required to keep holding the keytouch device through another variable delay of 1000-1500 after which the red keytouch cue disappeared then the NHP was trained to release its hand from the keytouch prompting the commence of the delay (1000ms) between sample and choice-phases. Then after the 1000 ms delay period another key-touch red dot cue, this time in the bottom of the screen, and the animal was required to hold the key-touch again. The NHP was again required to keep holding the key-touch device through a variable delay of 1000-1500 ms after which two choice stimuli (one an object image and the other a black circle, left-right randomized between trials) appeared. The NHP was required to keep holding the keytouch device through another variable delay of 1000-1500 ms after which the key-touch red dot cue disappeared cueing the animal that it could release holding the key-touch and make a choice to the touchscreen to either the object test-image stimulus or to the black circle. Just as in the human task in Figure 2, in ‘Match Trials’ the black circle was presented with an identical image to one of the preceding sample; in ‘Non-match Trials’ the black circle was presented with a novel stimulus not seen before. Animals were trained to touch the test-item if they remembered the test-item was a match but to touch the black circle if they thought the test-item was a non-match; accordingly the responses could be separated into hits, misses, correct rejections, and false alarms as indicted. Panel B: drawing of a lateral view of the NHP brain; the Utah array was placed above the principle sulcus (PS) and anterior to the arcuate sulcus (Arc); the enlarged insert (red square) shows photographic verification of the array’s position.

#### 2.1.3 Task stimuli and apparatus

The object recognition memory task was programmed using Turbo Pascal (Borland), run under DOS on a desktop PC. Visual stimuli used in the task were clip-art images in colour, which were presented on a 20.1” colour touch-sensitive screen (TFT LCD TS200H GNR). Those clip-art images used in the task were from a large pool of several thousand unique images. Each image subtended 5° of visual angle in width and 5° in height to the subject when presented on the screen. The sample image was always presented on the right top of the screen, positioned +9 ° horizontal and −9 ° vertical from the center of the screen. The test images were presented on the right bottom of the screen: one was positioned 0 ° horizontal and +3 ° vertical from the center of the screen; and the other one was positioned +17 ° horizontal and +3 ° vertical from the center of the screen. The background colour to the screen was white. In each session of recordings, images were randomly chosen from the pool without replacement and were not re-used on the other recording days.

The animal was seated in a primate chair (Rogue Research Inc.) in front of the touch screen with its head-fixated and whilst it performed the recognition memory task in a magnetic-shielded, and partially sound-attended, testing-box. A window in the front of the chair provided its access, both to the touch-screen itself and also to a touch-sensitive knob which we refer to as a ‘key-touch’ which was positioned low down in front of the touch screen; the animal had to steadily hold the keytouch at various times in each trial (to control for arm movement/position whilst it waited for stimuli, looked at stimuli, and waiting for a visual cue to release keytouch and touch the screen to make a choice). The distance between the monkey and touch screen was fixed at 50 cm enabling the animal to touch the screen easily. An infrared camera was used to monitor the general status of the monkey in the box. A peristaltic pump device located on top of the box fed smoothie reward through a tube and to a spout positioned in the vicinity of the animal’s mouth. Below the screen was also an automated lunch-box which contained the majority of the animal’s daily meal (wet mash and fruits and nuts etc.) and which opened immediately at the end of the task.

#### 2.1.4 Behavioural task

The task is a variant of the well established delayed-matching-to-sample (i.e. recognition-memory) paradigm in which a stimulus in a sample phase has to be judged as familiar or not in a subsequent choice phase after a short delay; the form of the task used is similar to that used by Basile and Hampton (2013) in that in the choice phase, to allow separation of hits, misses, false alarms, and correct rejections, one choice image is presented with one non-match button (black circle) such that the animal selects the choice image if it considers the choice image a match and selects black circle if it considers it is a non-match (see Fig.1A). In each trial, the animal initiated the task by holding the key-touch when cued to do so by a small red circular keytouch cue (located towards upper centre of screen; Fig. 1A) presented on the screen. The NHP was required to keep holding the keytouch device through a variable delay of 1000-1500 ms (if the keytouch hold was broken the trial aborted) after which a central sample (object image) appeared behind the red key-touch cue. The NHP was required to keep holding the keytouch device through another variable delay of 1000-1500 ms (again, if the keytouch hold was broken the trial aborted) after which the red keytouch cue disappeared which was the cue to the animal it could release its hand from the keytouch at which point the delay (1000ms) between sample and choice-phases began (the maximal time for releasing key-touch was 5000 ms else the trial aborted and the animal received a time-out for 10 s). Then after the 1000 ms delay period another red keytouch cue appeared, this time in the bottom of the screen (close to, and equidistant, from where the two choice items would appear) and animal was required to hold the key-touch again. The NHP was again required to keep holding the keytouch device through a variable delay of 1000-1500 ms (if the keytouch hold was broken the trial aborted) after which two choice stimuli appeared (one an object image and the other a black circle, left-right randomized between trials). The NHP was required to keep holding the keytouch device through another variable delay of 1000-1500 ms (again, if the keytouch hold was broken the trial aborted) after which the red keytouch cue disappeared which was the cue to the animal that it could release holding the keytouch and now make a choice to the touchscreen to either the object test-image stimulus or to the black circle stimulus. The object stimulus was either the identical stimulus to the sample seen earlier in the trial or it was not identical to the sample. The animal was rewarded by delivery of 10 ml of smoothie for touching the test-item image if it matched the sample image (these we refer to as ‘match trials’), or it was rewarded for selecting the standard ‘non-match button’ (i.e. the black circle) if the test-item was a non-match (these we refer to an ‘non-match trials’). After a correct response, the intertrial interval was 3 s. However, following any error trial (including both incorrect response and aborted trials), the intertrial interval was 10 s Accordingly, on *match trials* the animal could either make a correct response (‘hit’) or an incorrect response (‘miss’) whereas on *non-match trials* the animal could either make a correct response (‘correct rejection’) or an incorrect response (‘false alarm’); in this way the paradigm is similar to one previously used by Basile and Hampton (2013) but a key difference is that we did not restrict the stimulus set to just two stimuli as we used larger sets moreover we also varied the degree to which stimuli were either familiar or novel in the session. Specifically, in any given session 50% of the trials used trial-unique stimuli and 50% were ‘repeat’ stimuli used previously in the session (but not used in any previous session); in each session there were six repeat stimuli sorted into 3 pairs such that in each trial with repeat stimuli one pair was chosen at random and one member of the pair was randomly chosen to be the sample. Each session typically consisted of 150 trials so the six repeat stimuli got steadily more familiar across the session.

#### 2.1.5 Electrophysiological Recordings

Data were recorded simultaneously from the 32-electrodes in the microelectrode array implanted in dlPFC throughout the recognition memory task. We only considered the trial-unique stimuli in these analyses as we used the trial-unique stimuli in human TMS study. The NHP performed, on average, 35 trials (with trial-unique stimuli) per day; the total average session duration was 45 minutes. In this study we analysed data recorded over 29 sessions approximately 8-12 weeks after the array implantation. The NHP had learnt the task in full prior to array implantation. Neural signals from each microelectrode were amplified, digitized at 30 kHz using a 256-channel Cerebus^TM^ Neural Signal Processor (Blackrock Microsystems). The local field potential (LFP) was analyzed off-line using the FieldTrip toolbox (http://www.fieldtriptoolbox.org, details see Oostenveld et al., 2011) for MATLAB (R2017b, MathWorks), as well as custom scripts (available upon request from corresponding author).

#### 2.1.6 Signal analysis

We investigated the power of the induced LFP responses during the encoding (i.e. sample presentation) phase of the NHP recognition memory task. A notch filter, the second-order infinite impulse response (IIR), was applied to the raw data signals, to remove the power from line noise (i.e. 50 Hz) and its harmonics (i.e. 100, 150 Hz). Then the data signals were down sampled from 30 kHz to 1 kHz. To obtain the power spectrums of LFPs for each of the 32 channels, a time-frequency decomposition analysis based on Morlet wavelet transform was performed. The frequency range in LFP spectral analysis was chosen to start at 4 Hz (to exclude slow-wave LFP components) and terminate at 120 Hz, in steps of 1 Hz.

For the 32 channels in the dlPFC array, the induced power spectra from LFPs were limited to data segments which contained completed trials and inter-trial intervals (ITIs). In each trial the duration of sample image presentation was 1000-1500 ms while the NHP held the key-touch throughout to control for hand movement; our analyses of LFPs considered a point 1000 ms before stimulus onset until 1000 ms after stimulus onset were analyzed. A 1000 ms segment during the ITI was picked as the baseline for all the power spectra, and their root mean square (RMS) was divided from the raw data before the calculation of the spectrum using a wavelet transform. In the recognition memory task, LFPs in both the pre- and post-sample presentation periods (both within a time window of 1000 ms while the NHP held the keytouch) from completed correct trials (N = 646) and completed incorrect trials (N = 241) were selected and analyzed.

#### 2.1.7 Statistical methods

To test for statistical significance of differences of the induced LFP power spectra between pre- and post-sample presentations during the recognition memory task, we performed a nonparametric permutation test, with the median difference between the above two conditions (i.e. pre and post-sample) as our test statistic. The non-parametric permutation test is an assumption-free method without prescribing underlying distributions (Maris and Oostenveld, 2007). In each trial, the time window of pre- and post-sample condition for nonparametric permutation test was 800 ms (i.e. time zero was the stimulus onset), to avoid the spectral leakage of LFP power when calculated near the margin of a 1000 ms time-window. This nonparametric permutation test was performed separately for correct trials and incorrect trials. In each trial type, the null hypothesis of this test was assumed to be that there was no significant difference of modulation of LFP power when comparing the observed difference between the two conditions (i.e. pre- and post-sample) against a reference distribution of differences between the two randomly assigned conditions. The reference distribution was obtained by performing 10,000 permutations on trial labels to randomly distribute the two conditions at at each frequency and at each time point. The observed median differences between the two conditions were selected whose values were larger than the 97.5th percentile of the maximum or smaller than the 2.5th percentile of the minimal in this reference distribution (*p* < 0.05, two-tailed). Then the selected values were clustered in connected sets on the basis of temporal adjacency. We calculated cluster-level statistics by taking the sum of the median-difference values within a cluster and took the largest of the cluster-level statistics as the significant differences between the two conditions. This procedure is a two-tailed test with a global false positive rate of 5% and correction for the multiple comparisons across frequencies and time points (Maris and Oostenveld, 2007).

To test for statistical significance of differences of the induced LFP power spectra between correct trials (N=646) and incorrect trials (N=241) during sample presentations, we performed a nonparametric permutation test, with the median difference between the above two conditions (i.e. correct and incorrect conditions) as our test statistic. In each trial, the time window of correct and incorrect conditions for the permutation test was 0-400 ms (i.e. time zero was the stimulus onset) as the above nonparametric permutation tests confirmed a significant cluster appearing from the sample image onset until 300 ms in both correct and incorrect conditions. The null hypothesis of this test was assumed to be that there was no significant difference of modulation of LFP power when comparing the observed difference between the two conditions (i.e. correct and incorrect conditions) against a reference distribution of differences between the two randomly assigned conditions. The reference distribution was obtained by performing 10,000 permutations on trial labels to randomly distribute the two conditions at at each frequency and at each time point. The observed median differences between the two conditions were selected whose values were larger than the 97.5th percentile of the maximum or smaller than the 2.5th percentile of the minimum in this reference distribution (*p* < 0.05, two-tailed). Then the selected values were clustered in connected sets on the basis of temporal adjacency. We calculated cluster-level statistics by taking the sum of the median-difference values within a cluster and took the largest of the cluster-level statistics as the significant differences between the two conditions.

### 2.2 Experiment 2: Human TMS study

#### 2.2.1 Participants

27 participants (18 male, 9 female, age range 18-30 years) took part in the TMS study. Participants were fluent English speakers, right-handed, and had normal or corrected-to-normal vision. Prior to the study, all the participants provided written consent and went through safety screening check to make sure they had no history of previous or current neurological or psychiatric conditions and were not taking any psychoactive medication. All the participants received monetary compensation for their participation at a standard rate for volunteers for TMS studies in Oxford. This study was carried out with the approval of Medical Sciences Interdivisional Research Ethics Committee, University of Oxford.

#### 2.2.2 Stimulation Brain Sites

In this study, we investigated the differential effects of rTMS to dlPFC and dmPFC in recollection and familiarity in a recognition memory task designed to be similar to that used with NHP in Experiment 1. We chose left dlPFC which approximately corresponded to area BA46 or BA9/46 (referred to as BA 9/46) in light of Experiment 1, and we chose dmPFC along the midline as an area the literature led us to expect to provide a contrast with respect to consideration of contributions to recollection and familiarity, this more medial area corresponded to area BA 8 or BA9 (referred to as BA 8/9). A further control brain site was chosen to be vertex, which we had no reason to expect to be important for either memory process, and which was localized at center point of the head. The three targeted sites were measured using Beam F3 localization system (Beam et al., 2009). This system allows the measurement of the location of the F3 electrode position in the 10-20 EEG coordinate system, and takes into account individual skull variations. Based on this system, BA 9/46 was localized 1 cm caudal to F3. Both vertex and BA8/9 sit on the midline, in which vertex was localized at a site corresponding to location of electrode Cz (measured as half the distance between inion and nasion and intersecting with half the distance between the two aural tragus); and BA8/9 was localized 7cm rostral to the vertex and 6 cm medial to F3. The relative positions of three stimulation sites are illustrated in Fig. 4A. The participants were assigned to one of the three brain site groups to equalize numbers per group: dlPFC group (BA 9/46, 9 participants), dmPFC group (BA8/9, 9 participants) and control group (9 participants).

#### 2.2.3 Defining Participant’s Motor Threshold

Prior to the experiment, all the participants were invited to a taster session, in which the individual’s Resting Motor Threshold (RMT) was obtained for each participant. To measure RMT, stimulation was applied over the site of left primary motor cortex (M1) (localized 5 cm laterally and 5 cm rostrally to the vertex), where the largest twitch in right index finger of the participant was found. Site search was initially started at the lowest stimulator output. By gradually increasing output, the RMT of the participant was determined when 5 out of 10 TMS pulses caused a twitch in the right index finger (Rossini et al., 1994).

A stimulation intensity of 90% of RMT was used in the study. This intensity was within an appropriate range taking into account the average scalp-cortical surface distance between M1 and stimulation areas and a 2.8% reduction in stimulator output for every mm closer to the skull (Stokes, 2005; Stokes et al., 2013, 2007). The mean RMT of dlPFC group was 44.33 (SE = 3.61), and of dmPFC group was 51.33 (SE = 5.52), and of vertex group was 49.11 (SE = 7.39).

#### 2.2.4 Task Stimuli and Apparatus

The task was an object recognition memory task similar to the one used in NHPs, similarly programmed using Turbo Pascal (Borland), run under DOS on a desktop PC and presented on a 20.1” colour touchscreen (TFT LCD TS200H GNR). The objects images used in the task were clip-art images as in NHP study, but in order to increase difficulty (in light of a pilot study wherein performance was close to ceiling) the images were all converted to grey-scale and the contrast toned down in an attempt to make them harder to discriminate from each other. Additionally, the samples were presented in lists followed by lists of choice trials, again to reduce ceiling effects in the human version of the NHP task. Each image subtended 10° of visual angle in width and 10° in height to the participant sitting facing the screen. The sample image was always presented on the right top of the screen, positioned +12 ° horizontal and −12 ° vertical from the center of the screen. The test images were presented on the right bottom of the screen: one was positioned 0 ° horizontal and +5 ° vertical from the center of the screen; and the other one was positioned +23 ° horizontal and +5 ° vertical from the center of the screen. The background colour to the screen was white.

Participants sat with their eyes a distance of 25 cm from the screen, wearing earplugs in order to avoid audio disturbance of TMS pulses, resting their chins on a chin-rest and their foreheads on a head holder to stabilize their head position throughout the experiment. They were instructed to respond to items by touching them on the screen and gestured their confidence ratings using their right hands. For example, they indicate by raising fingers (1, 2 or 3) whether their opinion corresponding to their being somewhat confident (1), moderately confident (2), or absolutely confident (3) in their judgment as to whether they considered the test-item to be old (i.e. presented before in the preceding list as a sample) or new (not seen before in the preceding list as sample). None of the samples in this task were used in more than one list so all stimuli were trial-unique (and hence compared to the novel/trial-unique stimuli in the NHP task).

Transcranial Magnetic Stimulation (TMS) was carried out using a biphasic Magstim Super Rapid^2^ magnetic stimulator (Magstim, Dyfed, UK) with a double 70-mm figure-of-eight coil. The TMS coil was clamped in line with participants’ assigned brain sites and localized at a 45° angle off the midline with the handle pointing to the posterior direction throughout the experiment.

#### 2.2.5 Behavioural task

Prior to the experiment, participants were given instructions on how to perform the task, and how to indicate their confidence judgments (see above), and then they were introduced to the behavioural task in a short training/practice session (comprised of one list of 12 samples and then 12 test-item trials) of the task without stimulation to provide an opportunity to become familiarized with task procedure. We used lists in the human version to avoid ceiling effects as pilot investigations revealed the NHP task with single samples and single test-trials to be too easy for participants. The experimental session itself contained 3 sub-sessions, with each sub-session containing 15 blocks of trials. Participants took a 10-15 min break between each sub-session. Each block contained an encoding phase (i.e. a list of 12 sample images), and then a short delay, and then a test phase (i.e. a list of 12 test-item trials). The task structure is depicted, for one block, in Figure 2; during the encoding phase, 12 sample images were presented sequentially on the screen (individually for 480 ms), with inter-stimulus-interval as 350 ms. Participants were instructed to view and try to remember each sample image. After the sample phase (all 12 sample items) was completed, a blank screen was presented for 1 s (delay), and then the test phase commenced (all 12 test trials). Test trials were either ‘match trials’ or ‘non-match trials’. In each test trial, either an identical stimulus to one of the preceding 12 samples (i.e. ‘match trials’) was shown as a test image, or a novel and previously unseen image (i.e. ‘non-match trials’) was shown as a test image, and that test image appeared on the screen together with a black circle (the left/right position of the black circle and the test-trial image were randomized between trials). The match and non-match trials were also counter balanced (6 of each per test phase of 12 test-trials) and were put in a random order. Just as in the NHP version of the task (Experiment 1) participants were instructed to touch the test image if they thought it matched one of the 12 sample images, or touch the standard ‘non-match button’ (i.e. the black circle) if they thought the test image was not a match. Participants could touch the screen to make responses once the test image and black circle were on the screen, with a maximum responding duration set to 5000 ms (i.e. test trial ended if participants didn’t make responses within 5000 ms). After responding to the test image, participants were instructed to rate their confidence as to whether the test item was new or old using a scale of 1-3 by making three different movements with their fingers, corresponding to somewhat, moderate, and absolute confidence. Participants were further instructed to try to use the entire range of confidence responses as best they could and not simply select the extremes of confidence as defaults. The inter trial interval was 1000 ms.

**Figure 2.**
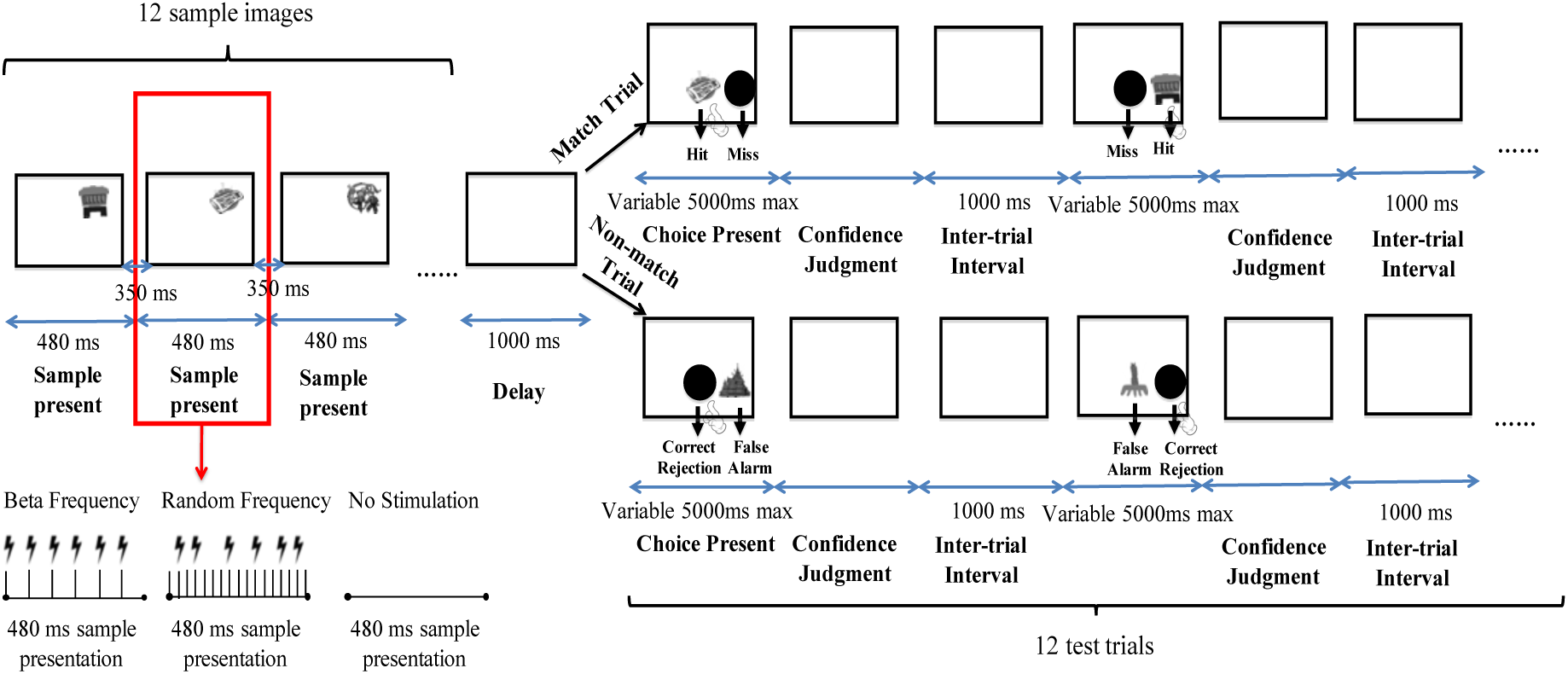
Human recognition memory task structure. The figure depicts one block in the TMS study. Twelve sample object images were sequentially presented for 480 ms with an inter-stimuli-interval of 350 ms (the figure shows the procedure for one of these 12). Six rTMS pulses (at beta frequency or at random frequency) or no stimulation were given during the 480ms sample presentation for 4 out of the 12 samples in the list. Then after a delay of 1000 ms, 12 test trials followed (the figure shows the procedure for one of these 12). Just as in the NHP task in Figure 1, in ‘Match Trials’ the black circle was presented with an identical image to one of the preceding sample; in ‘Non-match Trials’ the black circle was presented with a novel stimulus not seen before. Participants were told to touch the test-item if they remembered the test-item was a match but to touch the black circle if they thought the test-item was a non-match; accordingly the responses could be separated into hits, misses, correct rejections, and false alarms as indicted. Each test-trial in this human version of the task lasted maximally 5000 ms and ended with a confidence judgment indicated by the participant by their making a finger gesture as described in the main text. The inter test-trial interval was 1000 ms.

#### 2.2.6 Repetitive TMS Protocol

As our local field potential (LFP) investigations in the NHP (Experiment 1) revealed heightened beta power in dlPFC when macaques viewed and encoded sample images in a similar recognition memory task, the extent of which was associated with success or error in subsequent macaque recognition memory performance, rTMS stimulation in this human study was targeted during the encoding phase of the memory task so as to assess whether that activity in that region at that time may be necessary for successful encoding and subsequent recognition in humans.

In line with the safety guidelines for TMS by Rossi et al. (2009) to avoid seizure risks caused by too much sequential high-frequency stimulation, a restriction was implemented that there were at least 2 non-stimulated images in between every stimulated image in each sample list of 12 items. Therefore, we determined all the possible combinations wherein 4 out of 12 sample images might be targeted, with at least 2 non-targeted intervening samples, and each block’s sample phase took one of those schedules selected randomly each time. During beta-stimulation condition blocks, each of the four rTMS targeted samples in the list (each with a duration of 480 ms as detailed above) were targeted with the delivery of 6 pulses every 80 ms (hence at 12.5 Hz and within the range of beta frequency); while in the random-frequency stimulation blocks, the same total number of pulses (6) were delivered less regularly over the same period of presentation time (specifically, the 480ms sample presentation epoch was divided into 30ms intervals and 6 of those, with the constraint that no two consecutive 30ms intervals could be chosen, were chosen at random to trigger a TMS pulse delivery). All these considerations of TMS frequency ensured we always kept well within the known safe ranges of TMS stimulation at high frequency and over extended durations of stimulation in line with the safety guidance of Rossi et al. (2009). Each sub-session contained 15 blocks, with each of the three ‘types’ of stimulation (i.e. beta-stimulation, random-stimulation, and no-stimulation) occurring 5 times per sub-session, randomly ordered. Each participant received the same sets of stimuli and same stimulation order. As there were 3 sub-sessions in total each stimulation condition was repeated 15 times per participant and so participants performed a total of 540 trials (i.e. 3 sub-sessions x 15 blocks per sub-session x 12 trials per block). As we were interested in the effects of stimulated sample items on the subsequent memory performance, in each participant, 126 trials in beta-stimulation condition (i.e. excluding unstimulated old items appearing in the test phase), 123 trials in random-stimulation (i.e. excluding unstimulated old items appearing in the test phase) and 180 trials in no-stimulation (i.e. 15 blocks per sub-session x 12 trials per block) were used for our between-block analyses.

#### 2.2.7 Data analysis

Behavioural effects of TMS were evaluated based on dual-process signal detection (DPSD) theory model. According to this model, both familiarity and recollection contribute to recognition memory (Yonelinas, 2002, 2001). A recollection index (R) and familiarity index (F) were extracted by fitting the model to data using a standard approach that minimized the squared difference between the observed and predicted data in each confidence rating bin. Specifically, participants were asked to express their confidence in each of their decision using a 6-point scale, where 1 implies absolutely certain old, 2 for moderately certain old, 3 for slightly certain old, 4 for slightly certain new, 5 for moderately certain new, and 6 for absolutely certain new. The cumulative hit rate was then plotted against the cumulative false alarm rate to create the ROC plot of each participant. To be more specific, the first point at the left-hand side of the plot was the hit rate and false alarm rate at confidence level 6. After that, the second point was for the combination of confidence levels 6 and 5, the third point for confidence levels 6 to 4 and so on. The cumulative hit rate and false alarm rate of all confidence levels was constrained to 1.0. Accordingly, we had five coordinate points for each ROC plot.

The ROC data was analyzed using the DPSD model. This model assumes recognition memory to be contributed towards by two distinct processes referred to as recollection and familiarity, and the cumulative proportions of the target-item and lure-item trials in each confidence level (CL_i_) are given by (Koen et al., 2017):

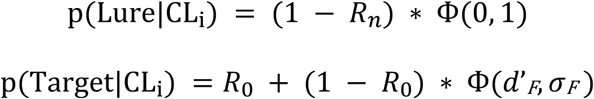

In the target and lure distribution (Ф is the cumulative response function), *R_0_* is a target threshold parameter being labeled recollection of old stimuli, and *R_n_* is a lure threshold parameter being labeled recollection of newness stimuli. In the lure item distribution, *R_n_* is a threshold parameter. If the strength of the lure item is beyond this threshold, it will always be classified as new. We have set *R_n_* to be 0 so the classification of lure item is always set as a standard normal Gaussian function with a mean of 0 and a unit standard deviation. Similarly, in the target item distribution, the *R_0_* parameter is the threshold for the classification of the target, which is an index of recollection, R. If the strength of a target item falls below *R_0_*, its classification would be governed by the familiarity component of the model which is a Gaussian distribution with the mean of *d’_F_* and standard deviation of σ*_F_*. The standard deviation σ*_F_* is commonly fixed to 1 in DPSD model, and the mean *d’_F_* is the index of familiarity, F. Based on the above assumptions, the algorithm of DPSD model is re-set as below:

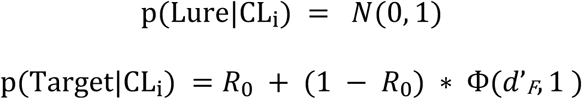

Each pair of the free parameters, *d’_F_* (F) and *R_0_* (R), defines a ROC curve. These parameters were obtained by minimizing the sum of squared residuals in both the abscissa and the ordinate of all the five points (that is, the sum of squared errors of prediction, or SSE, method for both axes). Based on the assumptions of the DPSD model, the range of R is between 0 and 1 while the range of F is between 0 and infinity (a theoretical maximum that is; practically it is much less, around 3-4 as a maximum). If the best fit of the *R* value turns out to be negative in the initial fitting of ROC and thus meaningless, we fixed it to be 0 and refit the model to get F. In each stimulation type and in each stimulation site condition, a ROC is plotted cumulatively for each confidence level by the proportion of correct ‘old’ judgements against the proportion of incorrect ‘old’ judgements, and both R and F in each stimulation type and stimulation site condition were extracted from DPSD model.

#### 2.2.8 Statistical methods

All the statistical analysis on human behavioral data was carried out using SPSS software (IBM). The above calculated indices (i.e. R and F) were subjected to 2×3×3 mixed model repeated measures ANOVAs with 3 levels of the within-group factor stimulation-type (beta-frequency, random-frequency, and no-stimulation) and with 3 levels of the between-subject factor stimulation-site (dlPFC, dmPFC, and vertex).

Accuracy was assessed for each human participant in each stimulation condition by calculating the area under each ROC curve (AUC). AUCs were calculated using the parameters from a fit of the DPSD model to the individual ROCs. Response time (RT) for correct responses were also evaluated for each participant under each stimulation condition. Then performance (i.e. accuracy or response time) was subjected to a 3×3 mixed model repeated measures ANOVAs with 3 levels of the within-subject factor stimulation-type (beta-frequency, random-frequency, and no-stimulation) and with 3 levels of the between-subject factor stimulation-site (dlPFC, dmPFC, and vertex).

## 3. Results

### 3.1 Experiment 1: NHP electrophysiological study

During the NHP recognition memory task, we focused only on the encoding (i.e. sample presentation) phase. A nonparametric permutation test as described earlier revealed a significant cluster ranging from 7-24 Hz starting after the sample image onset in macaque dlPFC in sample presentations preceding correct choices in subsequent choice-trials (two-tailed permutation test, *p* < 0.05) whereas a significant cluster ranging from 8-21 Hz was revealed starting after the sample image onset in macaque dlPFC in incorrect condition (two-tailed permutation test, *p* < 0.05). This post-sample alpha/beta power lasted roughly 300 ms in correct trials (Fig. 3A) and in incorrect trials (Fig. 3B). When comparing the difference between correct and error trials a further nonparametric permutation test (two-tailed permutation test, *p* < 0.05) revealed a cluster ranging from 10-15 Hz starting after the sample image onset in macaque dlPFC that was significantly higher in correct trials compared to incorrect trials. The analysis highlighter another small cluster ranging from 59-87 Hz (which was very brief, from 199 ms to 221 ms) found to be lower in beta power correct trials compared to incorrect trials. Therefore a stimulation frequency that was intermediate within this range (we chose 12.5 Hz, i.e. low beta) and that was, importantly, also well within the safety guidelines (with respect to duration and frequency considerations for repetitive stimulation) for humans (Rossi et al., 2009) was selected to be used in Experiment 2, our repetitive TMS (rTMS) study examining the effect of intervention at this frequency in the corresponding encoding phase of a homologous recognition memory task in humans.

**Figure 3.**
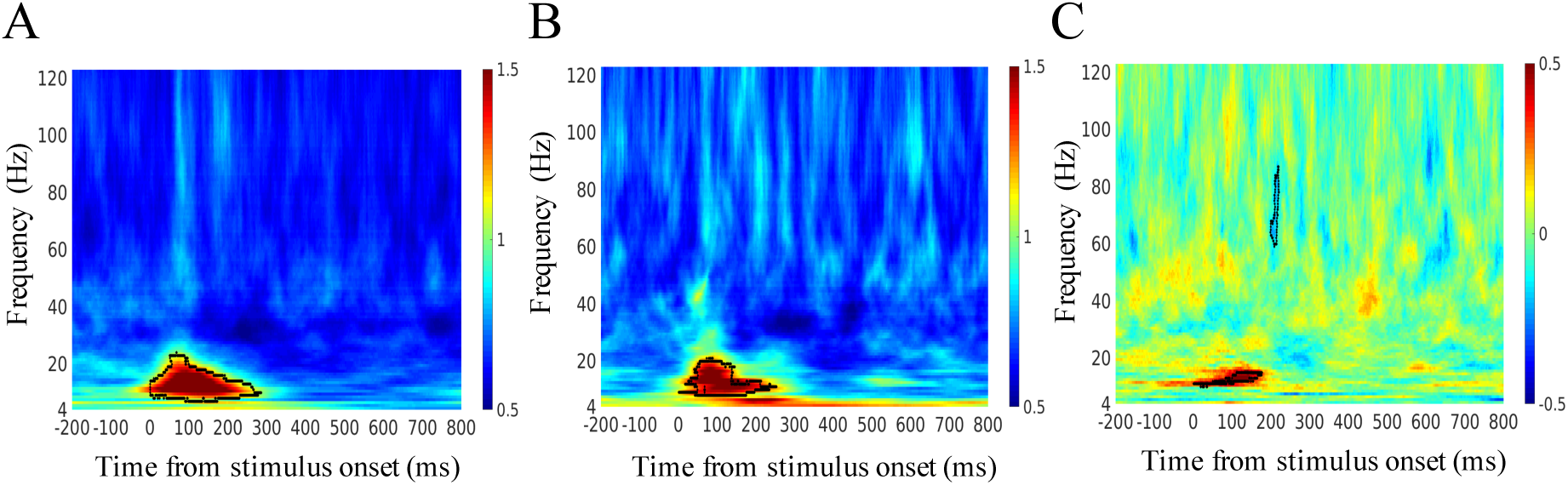
Time-frequency spectrogram for induced local field potential (LFP) power from dlPFC array in the sample presentation epoch in NHP in Expt.1. Panel A-B: the area of statistically significant differences of LFPs between pre- and post-sample periods in macaque dlPFC is highlighted by a black outline in trials that ended up being correct trials (A) and in trials that ended up being error trials (B). In both trials types we observed transient beta frequency power increase for around 300 ms from sample onset (indicated by zero ms on this figure). Panel C: the area of statistically significant higher LFPs in trials that ended up being correct trials compared to in trials that ended up being error trials in macaque dlPFC is highlighted by a black outline and includes low-beta frequencies in the 10-15 Hz range for around 200 ms from sample onset ms (indicated by zero ms on this figure); another area of statistically significant higher LFPs in error trials compared to correct trials is highlighted by another black outline and includes a very transient gamma peak in the 59-87 Hz range for around 20 ms. The data depicted in this figure is averaged across 32 electrodes in dlPFC, and across 29 daily sessions, and across 646 correct trials and 241 error trials.

### 3.2 Experiment 2: Human TMS study

To summarise the results (statistical analyses described in detail below) we observed that only rTMS at beta frequency to dlPFC drove down the recollection index, and only rTMS at beta frequency to dmPFC drove up the familiarity index.

### 3.3 Impact of TMS on accuracy and response time in the human recognition memory task

Participants were highly accurate in their recognition memory task; mean performance (calculated as AUCs) is shown in Table 1. A mixed-model repeated measures ANOVA, with stimulation-site as the between-subjects factor and stimulation-type as the within-subject factors, showed that neither the main effects (i.e. stimulation-site, stimulation-type) nor their interaction was significant (main effect of stimulation-site: *F*_(2,24)_ = 2.035, *p* = 0.153, *η^2^* = 0.145; main effect of stimulation-type: *F*_(2,48)_ = 1.252, *p* = 0.295, *η^2^* = 0.050; interaction: *F*_(4,48)_ = 0.529, *p* = 0.715, *η^2^* = 0.042) with respect to AUC.

**Table 1.**
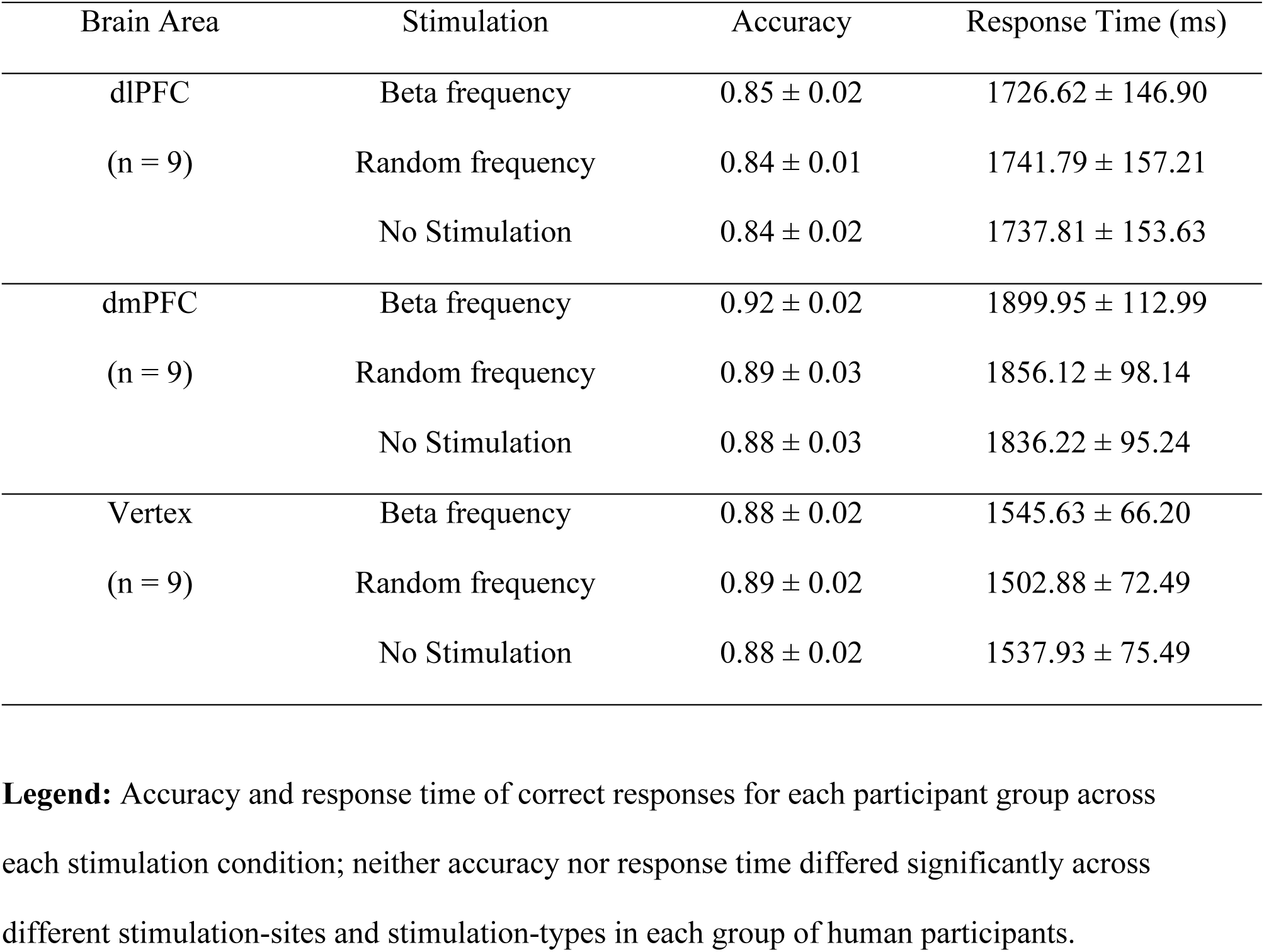
Descriptive statistics: accuracy (AUCs) and response time of correct responses for each participant group across each stimulation condition.

| Brain Area | Stimulation | Accuracy | Response Time (ms) |
| --- | --- | --- | --- |
| dlPFC<br>(n = 9) | Beta frequency | $0.85 \pm 0.02$ | $1726.62 \pm 146.90$ |
| | Random frequency | $0.84 \pm 0.01$ | $1741.79 \pm 157.21$ |
| | No Stimulation | $0.84 \pm 0.02$ | $1737.81 \pm 153.63$ |
| dmPFC<br>(n = 9) | Beta frequency | $0.92 \pm 0.02$ | $1899.95 \pm 112.99$ |
| | Random frequency | $0.89 \pm 0.03$ | $1856.12 \pm 98.14$ |
| | No Stimulation | $0.88 \pm 0.03$ | $1836.22 \pm 95.24$ |
| Vertex<br>(n = 9) | Beta frequency | $0.88 \pm 0.02$ | $1545.63 \pm 66.20$ |
| | Random frequency | $0.89 \pm 0.02$ | $1502.88 \pm 72.49$ |
| | No Stimulation | $0.88 \pm 0.02$ | $1537.93 \pm 75.49$ |

**Legend:** Accuracy and response time of correct responses for each participant group across each stimulation condition; neither accuracy nor response time differed significantly across different stimulation-sites and stimulation-types in each group of human participants.

Response times (RTs) were also calculated for each participant group in beta frequency, random frequency, and no-stimulation TMS conditions (Table 1). The aforementioned mixed-model repeated measures ANOVA applied to RT also showed neither the main effects (i.e. stimulation-site, stimulation-type) nor their interaction was significant (main effect of stimulation-site: *F*_(2,24)_ = 2.255, *p* = 0.127, *η^2^* = 0.158; main effect of stimulation-type: *F*_(2,48)_ = 1.184, *p* = 0.315, *η^2^* = 0.047; interaction: *F*_(4,48)_ = 1.387, *p* = 0.253, *η^2^* = 0.104).

### 3.4 Impact of TMS on recollection and familiarity in the human recognition memory task

The recollection index (R) and familiarity index (F) were calculated for each participant in each of the three stimulation types. A mixed-model repeated measures ANOVA was conducted on the indices data, with stimulation-site (3 levels: dlPFC, dmPFC, and vertex) as the between-subjects factor and stimulation-type (3 levels: beta-frequency, random-frequency, and no-stimulation) and index-type (2 levels: recollection-index, familiarity-index) as the two within-subject factors. A significant main effect of index type (*F*_(1,24)_ = 65.809, *p* < 0.001, *η^2^* = 0.733) was found, in addition to a significant two-way interaction between index-type and stimulation-type (*F*_(2,48)_ = 6.112, *p* = 0.008, *η^2^* = 0.203) with Greenhouse-Geisser correction due to a significant Mauchly’s test of sphericity. As tests of the linear trend component of the 3-way interaction between index type, stimulation type, and stimulation site were highly significant (*F*_(2,24)_ = 5.190, *p* = 0.013, *η^2^* = 0.302) we submitted the data to further scrutiny.

Therefore we next applied, repeated-measures ANOVAs to each stimulation-site independently, firstly considering the dlPFC site with the aforementioned index-type and stimulation-type within-subjects factors. We found a significant main effect of index-type (*F*_(1,8)_ = 82.060, *p* < 0.001, *η^2^* = .911) and a significant two-way interaction between index-type and stimulation-type (*F*(_2,16)_ = 3.984, *p* = 0.039, *η^2^* = 0.332), the latter prompted further scrutiny by a repeated-measures ANOVA on R alone, with stimulation type as the within-subjects factor, and this showed that although the main effect of stimulation was marginally significant (*F*_(2,16)_ = 4.432, *p* = 0.055, *η^2^* = 0.356; with Greenhouse-Geisser correction due to a significant Mauchly’s test of sphericity) the tests of within-subjects contrasts showed a highly significant effect of the linear component of stimulation type (*F*_(1,8)_ = 21.547, *p* = 0.002, *η^2^* = 0.729) which prompted further scrutiny by post-hoc pairwise t-tests with bonferroni correction. These revealed a significant suppression of the R index under beta stimulation compared with R under no-stimulation (*p* = 0.005), although no significant difference of the R index between beta stimulation and random stimulation was found (*p* = 0.272). Another parallel repeated-measures ANOVA, this time only considering the F index, with stimulation type as the within-subjects factor, showed that neither the main effect of stimulation type (*F*_(2,16)_ = 2.380, *p* = 0.125, *η^2^* = 0.229) nor the linear trend component (*F*_(1,8)_ = 2.680, *p* = 0.140, *η^2^* = 0.251) was significant.

Secondly, we ran another repeated-measures ANOVA this time considering just the dmPFC site, again with aforementioned index-type and stimulation-type as within-subjects factors. We found a significant main effect of index-type (*F*_(1,8)_ = 15.964, *p* = 0.004, *η^2^* = 0.666), a significant main effect of stimulation-type (*F*_(2,16)_ = 5.430, *p* = 0.016, *η^2^* = 0.404), and a significant two-way interaction between index-type and stimulation-type (*F*(_2,16)_ = 5.326, *p* = 0.017, *η^2^* = 0.400). The interaction prompted further scrutiny and so a repeated-measures ANOVA on the R index alone, with stimulation-type as the within-subjects factor, showed the main effect of stimulation type not to be significant (*F*_(2,16)_ = 2.185, *p* = 0.145, *η^2^* = 0.215). A parallel repeated-measures ANOVA on the F index alone, similarly with stimulation-type as the within-subjects factor, revealed a significant main effect of stimulation (*F*_(2,16)_ = 6.144, *p* = 0.010, *η^2^* = 0.434) and follow-up post-hoc pairwise t-tests with bonferroni correction showed that with respect to the dmPFC site a significant enhancement of the F index under beta stimulation compared to under no-stimulation (*p* = 0.012).

Thirdly, parallel analyses were conducted for the vertex site, again with index-type and stimulation-type as within-subject factors; here a significant main effect of index-type was found (*F*_(1,8)_ = 22.931, *p* = 0.001, *η^2^* = 0.741), but neither the main effect of stimulation-type nor the two-way interaction between index-type and stimulation-type were significant (main effect of stimulation-type: *F*_(2,16)_ = 0.160, *p* = 0.854, *η^2^* = 0.020; two-way interaction: *F*(_2,16)_ = 0.094, *p* = 0.911, *η^2^* = 0.012) so no follow-up analyses were required.

In summary, the between-block analyses above indicate (see Figure 4) that, when comparing ROC derived R and F indices only calculated from participants’ performance in choice trials that included samples that had appeared in the preceding list of samples and that had also been stimulated with rTMS (i.e. in the beta-stimulation and random-stimulation blocks) compared to all trials in the no-stimulation blocks (all choice trials could be included from the no-stimulation block because all were equivalent in that none of them had received rTMS), then the R index appears to be suppressed by beta stimulation in dlPFC (compared to non-stimulated trials in the no-stimulation block) but the F index appears to be enhanced by beta stimulation in dmPFC (compared to non-stimulated trials in the no-stimulation block). This highly intriguing result is limited in interpretational scope, however, because the above analyses clearly do not also show that beta-stimulation significantly suppresses the R index compared to random frequency stimulation rTMS, nor that beta-stimulation significantly enhances the F index compared to random frequency stimulation rTMS. To attempt to address this further we next considered what we refer to as ‘within-block’ analyses wherein we also calculate a within-block comparison of R and F indices for trials that used samples that had occurred in the sample phase that were targeted with rTMS (denoted +) versus trials that used samples that occurred in the sample phase that were not targeted with rTMS (denoted −); hence we refer to these four indices as R+, R−, F+, and F−. These indices cannot be calculated for the no-stimulation block as all trials are without rTMS so this analysis is restricted to beta-stimulation and random-stimulation blocks. Specifically, we investigated whether the within-block numerical difference between R+ and R− and also between F+ and F− differed between blocks, that is, whether these differences themselves depended upon whether samples were targeted with beta versus random rTMS. The aim was to further probe whether the beta-stimulation effect on R in dlPFC might, in this more sensitive within-block analysis, be somewhat more distinguished (i.e. statistically significantly as opposed to just numerically) from the random-frequency effect in dlPFC (and not just from the no-stimulation effect previously observed); similarly, we also aimed to probe if the beta-stimulation effect on the F index in dmPFC might be distinguished from the random-frequency effect upon the F index in dmPFC.

**Figure 4.**
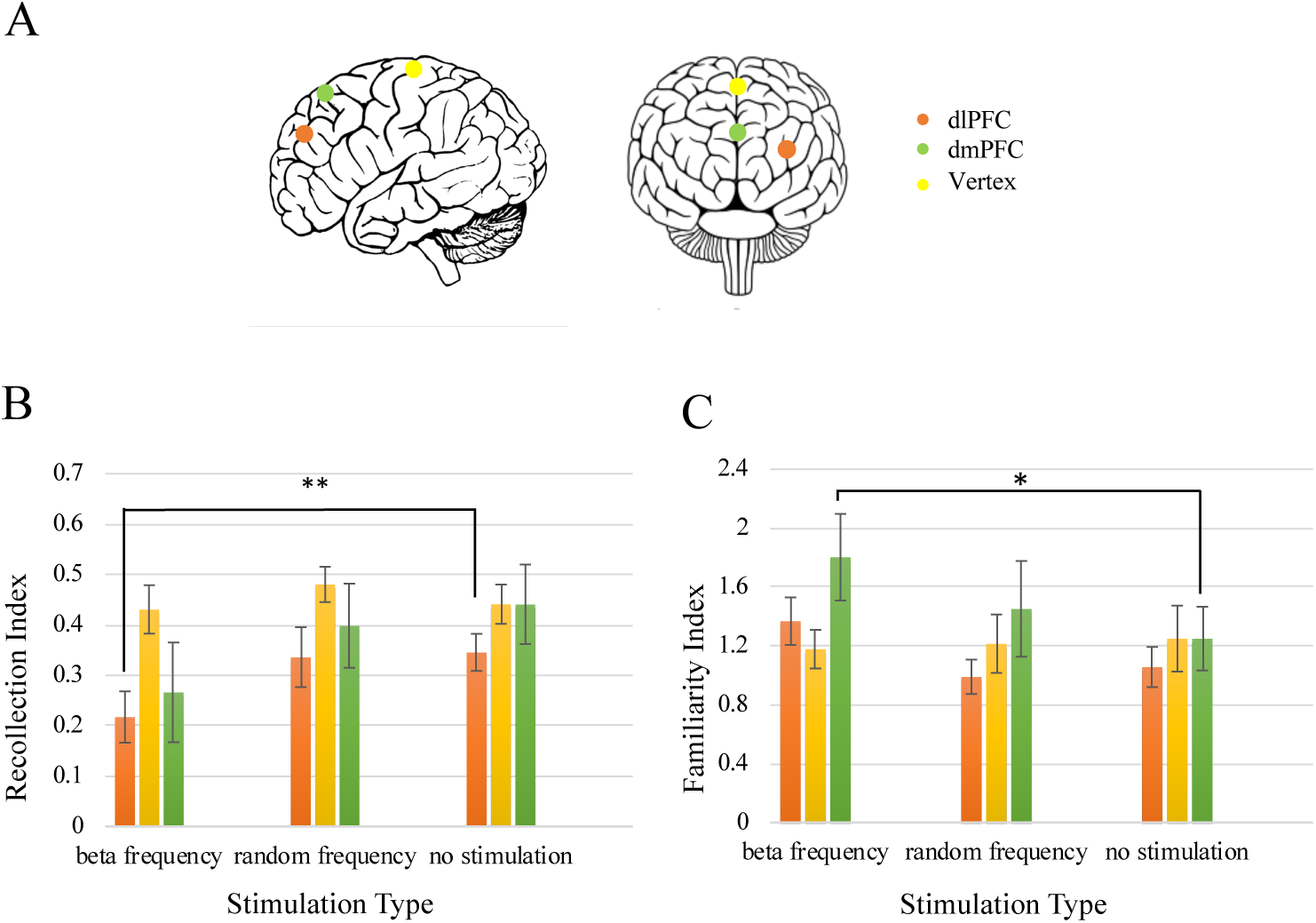
Stimulation sites for each group of human participants and effects of TMS upon recollection and familiarity indices in a between-block analysis in Expt. 2. Panel A: lateral and frontal view of the human brain. Human stimulation sites for rTMS are shown as dlPFC (orange), dmPFC (green) and vertex (yellow). Panel B-C: Bar graphs presenting the mean recollection index (Panel B) and familiarity index (Panel C) in dlPFC (orange bar), dmPFC (yellow bar) and vertex (green bar) under beta-frequency stimulation, random-frequency stimulation and no stimulation. Note that beta-frequency stimulation (but neither random-frequency stimulation nor no stimulation) suppressed recollection index in dlPFC, and beta-frequency stimulation (but neither random-frequency stimulation nor no stimulation) enhanced familiarity index in dmPFC. * indicates p < 0,05, and ** indicates p < 0.01.

Accordingly, a repeated measures ANOVA on the R+/R− difference observed in dlPFC with stimulation-type as a within-subjects factor (2 levels: beta-frequency block and random-frequency block) showed a significant main effect of stimulation-type (*F*_(1,8)_ = 6.508, *p* = 0.034, *η^2^* = 0.449) confirming a significant difference in this within-block analysis between the greater suppression of the R index by beta-frequency stimulation than by random-frequency stimulation (no significant difference by stimulus-type (*F*_(1,8)_ = 3.060, *p* = 0.118, *η^2^* = 0.277) was observed for the F+/F− statistic). In contrast, with respect to dmPFC, the repeated measures ANOVA on the R+/R− difference with stimulation-type as a within-subjects factor (2 levels: beta-frequency block and random-frequency block) did not show a main effect of stimulation-type (*F*_(1,8)_ = 0.143, *p* = 0.715, *η^2^* = 0.018) nor did a corresponding F+/F− analysis (*F*_(1,8)_ = 0.303, *p* = 0.597, *η^2^* = 0.037). Then, with respect to vertex, the repeated measures ANOVA on the R+/R− difference with stimulation-type as a within-subjects factor (2 levels: beta-frequency block and random-frequency block) did not show a main effect of stimulation-type (*F*_(1,8)_ = 0.996, *p* = 0.347, *η^2^* = 0.111) nor did a corresponding F+/F− analysis (*F*_(1,8)_ = 2.551, *p* = 0.149, *η^2^* = 0.242).

In summary the overall analyses (a combination of between-block and within-block approaches) provide mixed results and so must be interpreted cautiously; nonetheless the within-block analyses do at least support the notion that that the beta-frequency effect on the R index in dlPFC does not generalize to all rTMS stimulation parameters (as the random-frequency stimulation does not have the same effect) supporting the numerical trend seen in the between-block analyses and raising the possibility that it may be a frequency specific effect (how specific will need to be addressed by future studies); in contrast the beta-frequency effect on the F indices in dmPFC on the other hand appears to be less of a frequency specific effect as the difference between beta-frequency and random-frequency stimulation effects on the F indices did not attain statistical significance.

### 3.5 Impact of TMS on memory performance in match trials that containing samples that were either stimulated or not stimulated by rTMS

Finally, we returned, more directly, to the question of whether our rTMS protocols might be having effects that was clearly seen to be specific to choice trials that contained those particular samples (i.e. the 4 out of 12 in the preceding sample list) that were stimulated, or whether rTMS to those 4 samples might be having a broader effect upon performance across the block in general. We first calculated the hit rate for match trials that contained samples that had been targeted by rTMS in the sample list phase and compared that to the hit rate for match trials that did not contain samples targeted by rTMS in the sample list phase; the hit rate data was subjected to an ANOVA with the within-subjects factor ‘targeted’ (2 levels: rTMS-to-sample versus no rTMS-to-sample), the within-subjects factor stimulation-type (2 levels: beta-frequency block and random-frequency block), and with stimulation-site as the between-subjects factor (3 levels: dlPFC, dmPFC, and vertex). We found that there was a significant main effect of stimulation-type on this hit rate measure (*F*_(1,24)_ = 11.341, *p* = 0.003, *η^2^* = 0.321) and also a significant 3-way interaction between stimulation-type and targeted and stimulation-site (*F*_(2,24)_ = 4.264, *p* = 0.026, *η^2^* = 0.262) which prompted follow-up investigation of each stimulation-site separately which revealed that the main effect of the factor ‘target’ was not significant in any of the three regions (dlPFC: *F*_(1,8)_ = 1.156, *p* = 0.314, *η^2^* = 0.126; dmPFC: *F*_(1,8)_ = 0.106, *p* = 0.753, *η^2^* = 0.013; vertex: *F*_(1,8)_ = 0.277, *p* = 0.613, *η^2^* = 0.034) nor were interactions with stimulus-type (dlPFC: *F*_(1,8)_ = 2.767, *p* = 0.135, *η^2^* = 0.257; dmPFC: *F*_(1,8)_ = 0.242, *p* = 0.636, *η^2^* = 0.029; vertex: *F*_(1,8)_ = 4.679, *p* = 0.062, *η^2^* = 0.369). These analyses suggest that to some degree rTMS to just 4 our of 12 samples in the list had a broader effect on performance than to those targeted samples alone; this is to be expected as interference with sample encoding may lead to some broader confusion between presented and non-presented samples at choice. On the other hand our earlier within-block analyses reveal that R measures are different for R+ and R− conditions which implies that any effect is not equal to all choice trials. Likely there is both a degree of sample-focused rTMS effect and a degree of diffuse/spread of rTMS effect. Future studies that investigate the persisting effects of rTMS ideally need to also employ simultaneous recordings of EGG or neuronal responses so to correlate physiological measures with behavioural measures that outlast rTMS presentation.

## 4. Discussion

In this study, we first observed in a NHP what frequency-specific and performance related LFP power modulations occurred in dlPFC during sample image presentation in an object recognition/working memory task. An early transient peak in low-beta power was seen during sample presentation for 300 ms, moreover we observed a significant difference between its power during sample presentations in trials that ended up being correct recognition trials compared to in sample presentations that ended up being incorrect trials; specifically, the 10-15 Hz peak in low-beta power was significantly greater in the former than in the latter suggesting that the neuronal activity during sample presentation was indeed related to behavioural performance (i.e. subsequent recognition being correct versus incorrect). The significant brief bursts of beta power observed in our LFP signal between correct and error trials had a very early onset and only lasted around 200 ms. Previous electrophysiological investigation in NHPs have also found early involvement of dlPFC in the representation of salient stimuli, for example visual responses in the population peristimulus time histogram were observed with a latency as early as 42 ms and linked to bottom-up visual attention (Katsuki and Constantinidis, 2012). Early signals with a somewhat longer latency (∼120ms) observed in dlPFC have also been linked with top-down visual attention (Buschman and Miller, 2007). These aforementioned latencies are all well within the time-frame of the heightened beta power observed in our LFP recordings and so it is possible that the early signal we observed in dlPFC reflects both visual and attentional processes that benefited sample encoding and subsequent memory performance of NHPs in accordance with our observations of a behavioural association of signal magnitude with subsequent correct versus incorrect recognition. These LFP observations are limited with respect that to-date the data are from a single NHP providing information about task parameters to be used in a more comprehensive in-depth NHP study in progress but the link to behaviour gave us confidence to use them to inspire investigation of causality in a human TMS paradigm using a similar task.

Our initial hypotheses for the human TMS study were also inspired by the neural de-synchronization hypothesis, based on information theory, that was previously proposed (Simon Hanslmayr et al., 2012) to account for a *decrease* of alpha/beta LFP power in frontal and parietal cortex correlating with *successful* memory encoding and retrieval processes across some human studies (Hanslmayr et al., 2011, 2009; Khader and Rösler, 2011; Waldhauser et al., 2012). Specifically, power decreases in the alpha/beta frequency band are presumed to reflect a de-synchronization of local neural assembles. A rationale, based on information theory, is that synchronization of neural firings reduces the richness of information transfer which results in information redundancy; in contrast, a de-synchronization of neural firing patterns leads to more potential for carrying information to improve the efficiency of neural communications (Simon Hanslmayr et al., 2012). The de-synchronization hypothesis, applied to human recognition memory and our paradigm, maintains that a decrease of alpha/beta power may be associated with more efficient encoding, which helps contribute to successful memory retrieval subsequently (Simon Hanslmayr et al., 2012; Holmes et al., 2018). Therefore we first hypothesized that in our human investigations of a similar task to that performed by the NHP (albeit with lists of samples followed by lists of choice trials to increase memory demands so to avoid ceiling effects) that interfering in efficient beta desynchronization would be deleterious to successful encoding and hence recognition performance. To test this causal hypothesis we targeted rTMS stimulation at a beta frequency, chosen to be intermediate within the significant task-relevant low-beta frequency range found in our NHP observations (namely 12.5 Hz), and targeted to the homologous brain region (dlPFC) in human participants to that we recorded form in the NHP, all while they performed a similar task. We reasoned that artificial entrainment of low-beta frequency activity in dlPFC through 12.5 Hz low-beta rTMS throughout the 480ms of sample presentation might impair subsequent recognition by preventing effective beta desynchronization linked to effective sample encoding; therefore as a control rTMS stimulation protocol we also employed a random-frequency rTMS stimulation (i.e., with same total number of pulses over same total duration, but with each burst occurring pseudorandomly in time) presumed less likely to be deleterious to efficient and effective beta de-synchronization. We also targeted both rTMS stimulation protocols to both an active comparison site (dmPFC) given previously observed differences in relationship of these areas to recollective and familiarity processes in neuroimaging studies, and also a passive control site (vertex, which we had no reason to think was implicated in recognition memory).

We indeed found functional dissociations between the effects of short bursts of 12.5 Hz low-beta frequency rTMS stimulation (i.e. six pulses lasting 480ms, lasting the entire the duration of the sample presentation) to left dlPFC and left dmPFC. Only 12.5 Hz rTMS to dlPFC during sample presentations in the beta-stimulation block reduced behavioural indices of recollection derived from ROC/DPSD (see Methods) compared to performance in the no-stimulation block; whereas only 12.5 Hz rTMS to dmPFC in the beta-stimulation block enhanced similarly derived behavioural indices of familiarity compared to performance in the no-stimulation block (Figure 4 panel B and panel C). Short bursts of random frequency rTMS had no effect on either recollection or familiarity over either prefrontal regions, nor over the vertex. While intriguing these dissociations of beta-frequency effects between brain regions are limited with respect to claims about frequency specificity of the effects because the effects of beta-frequency stimulation, while differing from no-stimulation control, were not also significantly different from the random-frequency stimulation comparisons in this between-block analysis. To address this further we also conducted within-block analyses that applied ROC/DPSD separately to trials in each block that used samples that were targeted with rTMS and trials in each block that did not use samples targeted with rTMS; this ‘within-block’ approach was applied to beta-frequency blocks and random-frequency blocks wherein this distinction could be made. The within-block analyses revealed evidence that the effect of 12.5 Hz beta-stimulation to dlPFC on R index was significantly different from the effects of random-frequency stimulation to the same site on that index, but that the 12.5 Hz beta-frequency stimulation effect to dmPFC on the F index was not significantly different from the random-frequency stimulation effect on the same site on that index (Figure 5 panel A and panel B). Overall, our hypothesis as to the deleterious causal effects of low-beta frequency stimulation to human dlPFC sustained throughout the sample period was supported.

**Figure 5.**
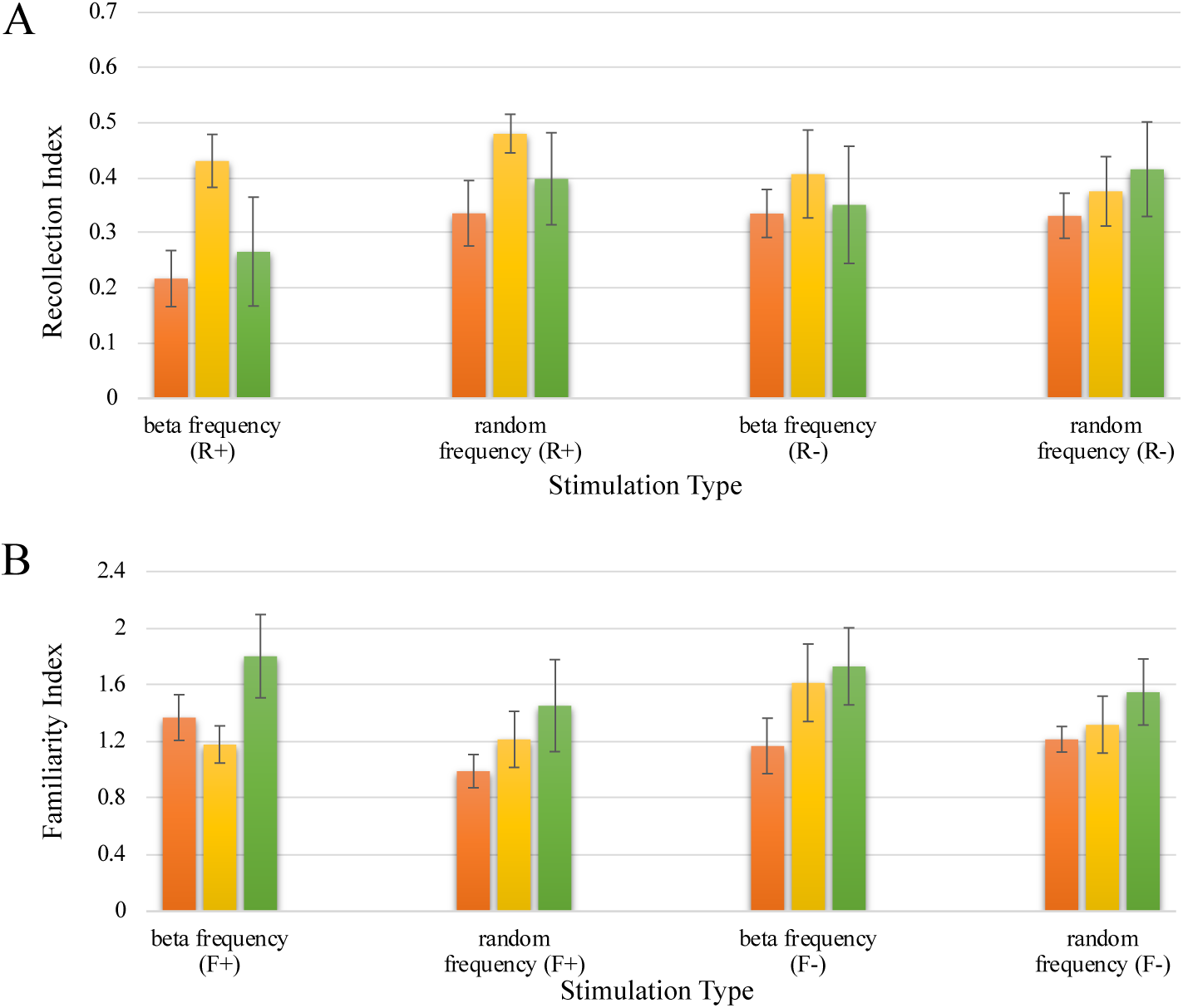
Stimulation sites for each group of human participants and effects of TMS upon recollection and familiarity indices in a within-block analysis in Expt. 2. Panel A-B: Bar graphs presenting the mean recollection index (Panel A) and familiarity index (Panel B) in dlPFC (orange bar), dmPFC (yellow bar) and vertex (green bar) under beta-frequency stimulation with samples targeted with rTMS (+) or without rTMS (-), and random-frequency stimulation with samples targeted with rTMS (+) or without rTMS (-). Accordingly, the R and F indices were labeled as R+, R−, F+, and F−. Our intended analyses compared the R+/R− and F+/F− difference scores and confirmed that these difference scores were significantly greater for R index dlPFC for beta-stimulation than for random-stimulation, but were not significantly greater for F in dmPFC for beta-stimulation than random-stimulation.

Another issue we considered was whether the rTMS effects we observed were specific to disrupted encoding of those particular samples viewed during rTMS stimulation or whether the causal effect upon subsequent recognition were attributable in part to a spread of effect, for example to all samples in the list given that it is known that rTMS effects can outlast the duration of stimulation (Hallett, 2007). Another possible explanation for general effect is that disrupted sample encoding may lead to a general confusion in evaluating familiarity or recollection of stimuli at choice. As we did not have EEG we sought behavioural evidence to speak to this issue. Firstly we note that our within-block analyses point to some specificity of rTMS effect (as ROC indices differed depending upon whether choice trials contained targeted samples or non-targeted samples), however when we analyzed the proportion of trials at choice that were correct (i.e. ‘hits’) for trials that used samples targeted with rTMS compared to those that did not we found no significant difference which suggests a spread of effect.

As no online EEG recording was carried out in our human’s task, this hypothesis of a de-synchronization effect of random-stimulation versus beta-frequency stimulations during encoding remains tentative and needs to be validated in future human research (or indeed in NHPs performing the same task wherein, as we showed, electrodes can be placed directly in the relevant regions, so that in future studies recording can be made in and around the regions that are stimulated). Previous studies have however combined EEG and TMS study, and have provided stimulation over left dlPFC (BA 9) using rTMS at a particular beta frequency (18.7 Hz), but not at other lower frequencies (6.8 Hz and 10.7 Hz), also impaired recall processes (Hanslmayr et al., 2014). This gives support to hence our external source of beta frequency entrainment from rTMS over the same region may have impaired recognition memory performance by lessening natural processes of de-synchronization as this hypothesis predicts. Holmes et al. (2018) also showed a dissociation between LFP beta power and mnemonic tuning and whilst in a spatial task this is also consistent with de-synchronization aiding memory stability (Holmes et al., 2018). Neural attractor network modeling (Compte et al., 2003), in the context of delayed response task performance in NHPs, also suggests mechanisms by which a drop in LFP power may relate to memory processing; asynchronous networks can maintain memory coding for longer durations whereas synchronization leads to an overall instabilities incompatible with stable memory-coding states. According to these models too, frontal memory networks desynchronize to preserve memory states.

The neural communication through coherence hypothesis (Fries, 2005) suggests that synchronized neural oscillations within a particular brain region or between distant brain areas provides important mechanisms for mediating cognition. This theory captures the general concept of interaction between spikes and oscillatory LFPs, and brain areas fluctuating between frequencies and/or phase of oscillations with respect to each other in order to ‘*tune-in*’ (Fell and Axmacher, 2011; Fries, 2015). Inter-area phase synchronization may have other effects too: e.g. facilitating timing of synaptic inputs onto common targets, or inducing spike timing-dependent plasticity of synaptic connections between regions facilitating communication (Fell and Axmacher, 2011). In a review focusing on frontal-temporal interactions in recollection/familiarity, a “multi-effect multi-nuclei” model was proposed, in which inputs to PFC directly from hippocampus, and indirectly from hippocampus *via* the anterior and midline thalamic nuclei, are related to recollection process, while connections to PFC from perirhinal cortex and mediodorsal thalamic nucleus are related to familiarity process (Aggleton et al., 2011). Indeed, a three-circuit model of neural oscillations between PFC and MTL *via* thalamus has been proposed (Ketz et al., 2015). The first theta-circuit model emphasized interaction between PFC and hippocampus through theta oscillations (4-8 Hz) to support core functions of the hippocampus in successful memory encoding and retrieval processes (Ketz et al., 2015). The human neuropsychology and neuroimaging literature highlight the involvement of the hippocampus (Bowles et al., 2010; Eichenbaum et al., 2007; Ranganath et al., 2004; Skinner and Fernandes, 2007; Yonelinas et al., 2002) and PFC (Wheeler and Stuss, 2003) in recollection, hence theta oscillations within a PFC-hippocampus network may be essential for recollection process. In our TMS study then, the external source of beta frequency from rTMS over dlPFC may disrupt the communication between PFC and hippocampus mediated by theta-oscillation, which may be a plausible mechanism by which our rTMS effected a decrease of recollection process. The theta-circuit model of dlPFC and hippocampus interactions in recollection processes will require further investigations and will shed light on the systems-wide oscillatory mechanisms underlying recognition memory. Recording simultaneously from multiple areas and multiple cortical layers in NHPs will further elucidate these issues.

Other studies have also applied the information theory by de-synchronization hypothesis to investigate voluntary memory suppression. A combined fMRI-EEG-TMS study has shown that an increased BOLD signal in left dlPFC (BA 9) was associated with decreased neural synchrony at beta frequency (11-18 Hz) in a directed forgetting task; moreover, stimulation over dlPFC using a slow train of 1 Hz rTMS pulses in the same task boosted cued forgetting-behaviour and induced a further reduction in neural synchrony at 13 Hz (S. Hanslmayr et al., 2012). This study also suggests a causal role for beta desynchrony in dlPFC, this time in a directed forgetting paradigm and also suggests a key role of dlPFC in memory control operates *via* downregulating neural synchrony at beta frequency (S. Hanslmayr et al., 2012). Another fMRI study has shown that voluntary memory suppression was associated with an increased activation of dlPFC and a decreased activation of hippocampus (Anderson et al., 2004), which suggests an interaction between dlPFC and hippocampus in regulating memory suppression.

In our human TMS study, familiarity was enhanced in association with rTMS delivered at beta frequency over dmPFC. This result is in line with human neuroimaging meta-analyses suggesting that more activation peaks related to familiarity processing in PFC were found in dmPFC (BA area 8 and 9) (Scalici et al., 2017). Our study not only shows a causal involvement of dmPFC in familiarity-based recognition, but suggests that the mechanism driving familiarity may operate through a mechanism involving neural synchrony at beta frequency. However, unlike for dlPFC, we didn’t detect a difference between beta-stimulation and random-stimulation over human dmPFC in familiarity, so our results indicate a potential mechanism of beta coherence in dmPFC modulating familiarity but they don’t rule out modulation effects of other frequency. In terms of potential mechanistic explanations for this finding, a thalamo-cortical circuit (involving entorhinal, parahippocampal, and perirhinal cortical areas connected via the medial dorsal thalamic nucleus to PFC) may synchronise within the beta frequency range and it has been proposed that this beta-circuit model may underlie familiarity judgment processes (Ketz et al., 2015). Indeed, the functional role of rhinal cortex, especially perirhinal cortex, has long been associated more with familiarity-based recognition than recollection (Aggleton and Brown, 1999; Bowles et al., 2010; Norman and O’Reilly, 2003; Ranganath et al., 2004; Skinner and Fernandes, 2007). TMS was a viable methodology in our study due to dlPFC and dmPFC both being accessible on the lateral surface of the frontal lobes, but the TMS approach is not viable for targeting deep structures in the medial temporal lobe. Future work in NHPs will of course be able to target microstimulation in via the same electrodes that record LFPs in the medial temporal lobe and record, simultaneously, the effects on multiple other brain regions.

The current project has helped bridge the species divide in understanding the neural mechanisms supporting object recognition memory between macaques and humans. The site of human rTMS stimulation, the frequency, and the stage of stimulation in the task were all inspired directly by study our NHP observations. Our rTMS study found evidence for dissociable functional roles of beta frequency oscillations in different prefrontal regions. As no online EEG recordings was carried out in combination with our rTMS stimulation the proposed function of beta coherence or even beta oscillations in human prefrontal regions in recollection and familiarity processes needs to be further validated in future studies using, for example, a combined EEG-TMS or MEG-TMS methodology, and/or for a more detailed neuronal level approach, in NHPs using multi-area multi-electrode neuronal recording combined with intervention methodologies including focal ultrasound, reversible pharmacological inactivation, or direct microstimulation through implanted electrodes. Importantly, our study helps further bridge the species divide and confirms that human and NHP PFC show similarity in function and in underlying neural mechanisms supporting recognition memory performance.

## Author contributions

**Zhemeng Wu:** Conceptualization, Data curation, Formal analysis, Investigation; Methodology, Software, Validation, Visualization, Writing - Original draft, Writing – Reviewing and Editing. **Martina Kavanova:** Data curation, Formal analysis, Visualization. **Lydia Hickman:** Data curation, Formal analysis, Visualization. **Fiona Lin:** Data curation, Formal analysis, Visualization. **Erica Boschin:** Supervision. **Juan M. Galeazzi:** Supervision. **Lennart Verhagen:** Resources, Supervision. **Mark J. Buckley:** Conceptualization, Data curation, Formal analysis, Funding acquisition, Investigation; Methodology, Project administration, Software, Supervision, Validation, Visualization, Writing - Original draft, Writing – Reviewing and Editing.

## Acknowledgements

This work was supported by Wellcome Trust Strategic Award Grant (Ref: WT101092MA) and MRC Project Grant (Ref: MR/K005480/1). L.V. is supported by funding from the Wellcome Trust (WT100973AIA). We thank the team of expert animal technicians and veterinary staff and anaesthetists for their very high standards of animal care and husbandry throughout. We thank Carlos Pedreira and Martin O’Neill for NHP support in setting up the electrophysiology lab. We thank Olof van der Werf and Alberto Lazari for T.M.S. support and we thank the Oxford Centre for Human Brain Activity (OHBA) and Matthew Rushworth for access to TMS facilities.

